# Genetic Effects of Welding Fumes on the progression of Neurodegenerative Diseases

**DOI:** 10.1101/480806

**Authors:** Humayan Kabir Rana, Mst. Rashida Akhtar, Md Bashir Ahmed, Pietro Lio’, Julian Quinn, Fazlul Huq, Mohammad Ali Moni

## Abstract

**Background:** Welding exposes different types of fumes, gases and radiant energy that can be potentially dangerous for unsafe welder’s health. Welding fumes (WFs) are a significant problem among all those exposed. WFs are a complex mixture of metallic oxides, silicates and fluorides that may result in different health effects. If a welder inhales such fumes in large quantities over a long period of time, there is a risk of various neurodegenerative diseases (NDGDs) development.

**Methods:** We developed quantitative frameworks to identify the genetic relationship of WFs and NDGDs. We analyzed Gene Expression microarray data from WFs exposed tissues and NDGDs including Parkinson’s disease (PD), Alzheimer’s disease (AD), Lou Gehrig’s disease (LGD), Epilepsy disease (ED), Multiple Sclerosis disease (MSD) datasets. We constructed disease-gene relationship networks and identified dysregulated pathways, ontological pathways and protein-protein interaction sub-network using multilayer network topology and neighborhood-based benchmarking.

**Results:** We observed that WFs shares 18, 16, 13, 19 and 19 differentially expressed genes with PD, AD, LGD, ED and MSD respectively. Gene expression dysregulation along with relationship networks, pathways and ontologic analysis showed that WFs are responsible for the progression of PD, AD, LGD, ED and MSD neurodegenerative diseases.

**Conclusion:** Our developed network-based approach to analysis and investigate the genetic effects of welding fumes on PD, AD, LGD, ED and MSD neurodegenerative diseases could be helpful to understand the causal influences of WF exposure for the progression of the NDGDs.

## 1. Introduction

Welding process is very dangerous because it exposes different types of fumes, gases and radiant energy. Welding fumes (WFs) are the most venturous partial among all welding exposers [1]. WFs are an intricate mixture of metallic oxides, silicates and fluorides include Beryllium, Aluminum, Cadmium, Chromium, Copper, Iron, Lead, Manganese, Magnesium, Nickel, Vanadium, and Zinc etc. [2]. If a welder inhales welding fumes in large quantities over a long period of time, this may convey various NDGDs [1, 3].

Neurodegenerative diseases (NDGDs) are a collective term for a heterogeneous group of disorders that are incurable and characterized by the progressive degeneration of the function and structure of the central nervous system [4]. NDGDs primarily attack the neurons of the central nervous system and progressively damage the function of them. Neurons are most vulnerable to injury and normally dont reproduce or replace themselves [5]. If neurons become damaged or die they cannot be replaced by medical treatments. So that NDGDs are very dangerous and currently they dont have any cure. We studied several NDGDs include PD, AD, LGD, ED and MSD to find the effects of welding fumes on them.

PD is the second-most neurologic disease that affects neural cells in the brain which produce dopamine in the substantia nigra [6]. There are several symptoms of PD include tremors, muscle rigidity, and changes in gait and speech. Welding fumes contain Manganese that can develop Parkinsons disease [7]. The AD is the most common type of incurable dementia that causes problems with progressive memory loss and other cognitive abilities. Existing medical treatments for AD produce only a modest improvement of symptoms but there is currently no cure [8]. Aluminum exposure to welding is a risk factor to produce AD. LGD also cognizant as Amyotrophic lateral sclerosis (ALS), is a neurodegenerative disease that progressively damages motor neurons and muscle atrophy controlling voluntary muscle movement. The initial symptoms of LGD are muscle weakness or stiffness, can bring death by progressive muscular paralysis and respiratory system failure within 2 to 5 years. US Food and Drug Administration (FDA) approved Riluzole and Edaravone drugs that could prolong LGD survival. But still, there is no effective cure or prevention for this devastating disease [9, 10]. ED is a heterogeneous group of neurodegenerative disorder that affects neural cells in the brain which are recognized by recurrent seizures or unusual behavior, awareness and sensations suffering over 60 million people in the world. AEDs are Current anti-epileptic drugs that can minimize symptoms but there is no permanent cure or prevention of ED [11]. MSD is a devastating neurodegenerative disorder that attacks the neurons of the central nervous system in the spinal cord and brain, on young adults most commonly [12]. The symptom of MSD can be included as muscle weakness, trouble with sensation, blindness in one eye or double vision. Medical treatments of MSD could prolong only survival but there is no permanent cure or prevention of MSD. Manganese exposure to welding is a main risk factor on the progression of LGD, ED and MSD [13].

Our study employed a systematic and quantitative approach to find the genetic effects of WFs on NDGDs. For these purposes, we studied several NDGDs including PD, AD, LGD, ED and MSD. To understand the effects of WFs on NDGDs, we examined gene expression dysregulation, disease association network, dysregulated pathway, gene expression ontology and protein-protein interaction. We also investigated the validation of our study by using the gold benchmark databases (dbGAP and OMIM).

## 2. Materials and Methods

### 2.1 Datasets employed in this study

To investigate the effects of WFs on NDGDs at the molecular level, we used gene expression microarray data. In this study, we used Gene Expression Omnibus from the National Center for Biotechnology Information (NCBI) (http://www.ncbi.nlm.nih.gov/geo/). We analyzed 6 different datasets for our study with accession numbers GSE62384, GSE19587, GSE28146, GSE833, GSE22779 and GSE38010 [14, 15, 16, 17, 18, 19]. The WFs dataset (GSE62384) is a result of gene expression analysis of fresh welding fumes influence on upper airway epithelial cells (RPMI 2650). This Data is collected from the people with spark-generated welding fumes at high (760 g/m3) and low (85 g/m3) concentrations. The donors inhaled welding fumes for 6 hours continuously, followed by zero hours or four hours post-exposure incubation. The PD dataset (GSE19587) is taken from 6 postmortem brains of PD patients and from 5 control brains. The AD dataset (GSE28146) is a microarray data on RNA from fresh frozen hippocampal tissue blocks that contain both white and gray matter, potentially obscuring region-specific changes. The LGD dataset (GSE833) is an Affymetrix Human Full Length HuGeneFL [Hu6800] Array. In this data, postmortem spinal cord grey matter from sporadic and familial ALGD patients are compared with controls. The ED dataset (GSE22779) is a gene expression profiles of 4 non-leukemic individuals (1 healthy and 3 with epilepsy) is generated from the mononuclear cells isolated from the peripheral blood samples before, and after 2, 6, and 24 hours of in-vivo glucocorticoid treatment. The MSD dataset (GSE38010) is a microarray data of multiple sclerosis (MS) patients brain lesions compared with control brain samples.

### 2.2 Overview of analytical approach

We used systematic and quantitative approach to identify the effect of WFs on the progression of the NDGDs using different sources of available microarray datasets. The graphical representation of this approach is shown in figure 1. This approach included gene expression, signaling pathway, Gene Ontology (GO) and protein-protein interaction analyses. This approach also used Gold benchmark data to verify the validity of our study.

**Figure 1:**
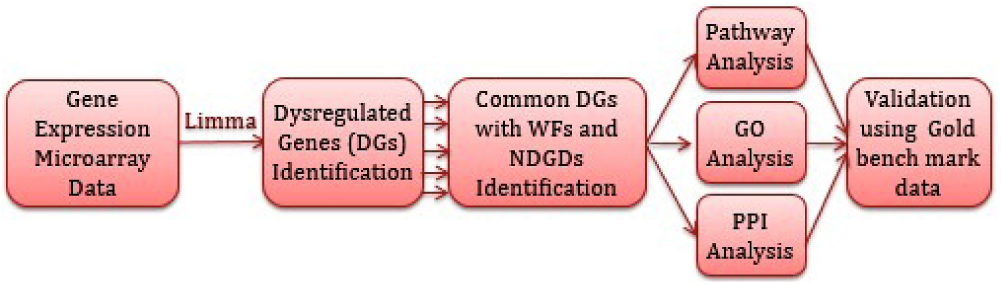
Flow-diagram of the analytical approach used in this study.

### 2.3 Analysis methods

Gene expression analysis using microarrays is a global and popular method to develop and refine the molecular determinants of human disorders that have proven to be a sensitive method [20]. We used these technologies to analyze the gene expression profiles of Parkinson’s disease (PD), Alzheimer’s disease (AD), Lou Gehrig’s disease (LGD), Epilepsy disease (ED) and Multiple Sclerosis disease (MSD) to find the effects of welding fumes on them. To uniform the mRNA expression data of different platforms and to avoid the problems of experimental systems, we normalized the gene expression data (disease state or control data) by using the Z-score transformation (*Z*_ij_) for each NDGD gene expression profile using

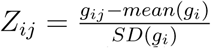

where *SD* implies the standard deviation, *g*_ij_ represents the value of the gene expression *i* in sample *j*. After this transformation we can directly compare of gene expression values of various diseases under different platforms. We applied two conditions for t-test statistic. We performed unpaired T-test to identify differentially expressed genes in patients over control data and selected significant genes. We have chosen a threshold of at least 1 *log*_2_ fold change and a *p*-value of <= 1*10^−2^.

We applied the topological methods and neighborhood based benchmark to find gene-disease associations. Gene-disease network (GDN) was constructed by using the gene-disease associations, where the nods in the network represent either gene or disease. This network can also be characterized as a bipartite graph. The diseases are connected in GDN when they share at least one or more significant differentially expressed genes. Let *D* is a specific set of diseases and *G* is a set of dysregulated genes, gene-disease associations attempt to find whether gene *g ∈ G* is associated with disease *d ∈ D*. If *G*_*i*_ and *G*_*j*_, the sets of significant dysregulated genes associated with diseases *D*_*i*_ and *D*_*j*_ respectively, then the number of shared dysregulated genes 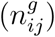 associated with both diseases *D*_*i*_ and *D*_*j*_ is as follows [20]:

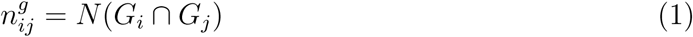

The common neighbours are the based on the Jaccard Coefficient method, where the edge prediction score for the node pair is as [20]:

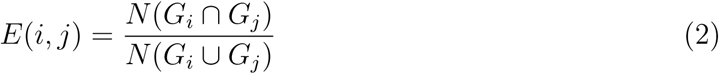

where *G* is the set of nodes and *E* is the set of all edges. We used R software packages “comoR” [21] and “POGO” [22] to cross check the genes-diseases associations.

To find molecular pathways of several NDGDs, we have analyzed pathway and gene ontology using Enrichr (https://amp.pharm.mssm.edu/Enrichr/), a comprehensive gene set enrichment analysis web-based tool. We used STRING (https://string-db.org.) for analyzing protein-protein interactions.

## 3. Results

### 3.1 Gene Expression Analysis

To investigate the effects of WFs on NDGDs, we analyzed the gene expressing microarray data from the National Center for Biotechnology Information (NCBI) (http://www.ncbi.nlm.nih.gov/geo/). We found that 903 genes were differentially expressed for WFs with adjusted *P <*= .01 and *|logFC|>*= 1. Among them, 392 and 511 were up and down regulated respectively. Similarly, our analysis identified the most significant differentially expressed genes for each NDGD after various steps of statistical analysis. We identified differentially expressed genes, 774 (263 up and 511 down) in PD, 565 (291 up and 274 down) in AD, 501 (296 up and 205 down) in LGD, 725 (350 up and down) in ED and 834 (455 up and 388 down) in MSD. The cross-comparative analysis was also performed to find the common differentially expressed genes between WFs and each NDGD. We observed that WFs shares 18, 16, 13, 19 and 19 differentially expressed genes with PD, AD, LGD, ED and MSD respectively. To find the significant associations among these NDGDs with WFs, we built two separate disease relationships networks for up and down-regulated genes, centered on the WFs as shown in figure 2 and 3. Two diseases are associated with each if there exist one or more common genes in between these diseases [23]. Noticeably, 2 significant genes, N4BD2L2 and NAAA are commonly differentially expressed among WFs, LGD and WD; one gene DAAM1 is commonly dysregulated among WFs, ED and MSD.

**Figure 3:**
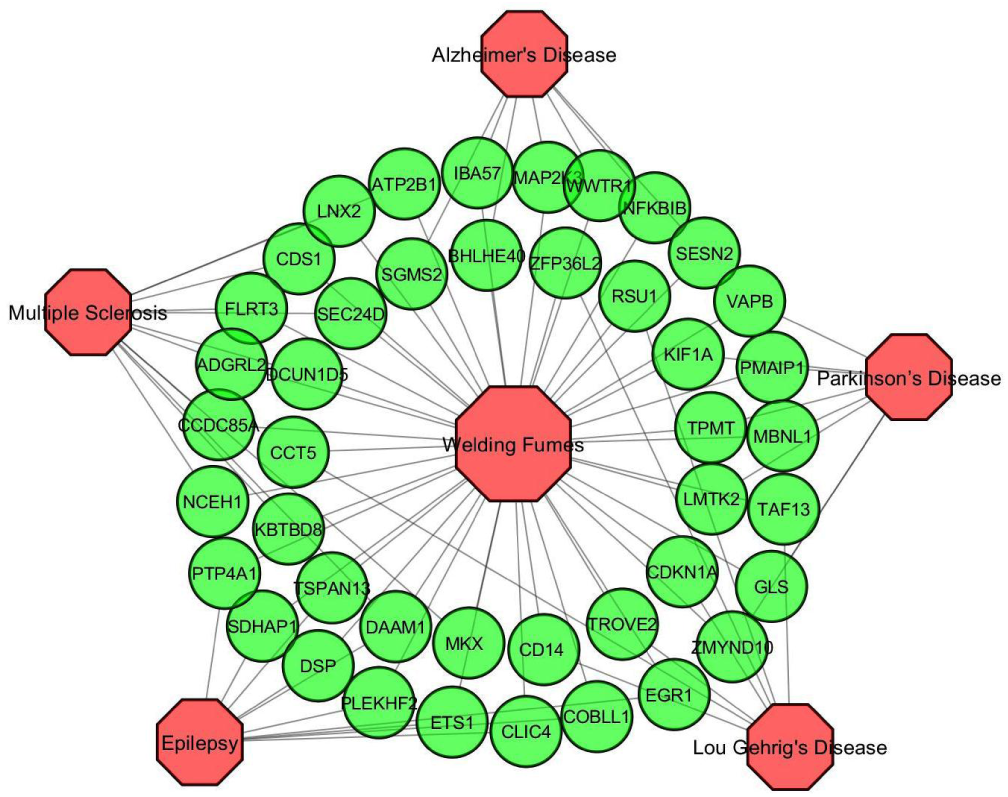
Disease network of Welding fumes (WFs) with Parkinson’s disease (PD), Alzheimer’s disease (AD), Lou Gehrig’s disease (LGD), Epilepsy disease (ED) and Multiple Sclerosis disease (MSD). Red colored octagon-shaped nodes represent different categories of disease, and round-shaped green colored nodes represent commonly up-regulated genes for WFs with the other neurodegenerative disorders. A link is placed between a disorder and a disease gene if mutations in that gene lead to the specific disorder.

**Figure 2:**
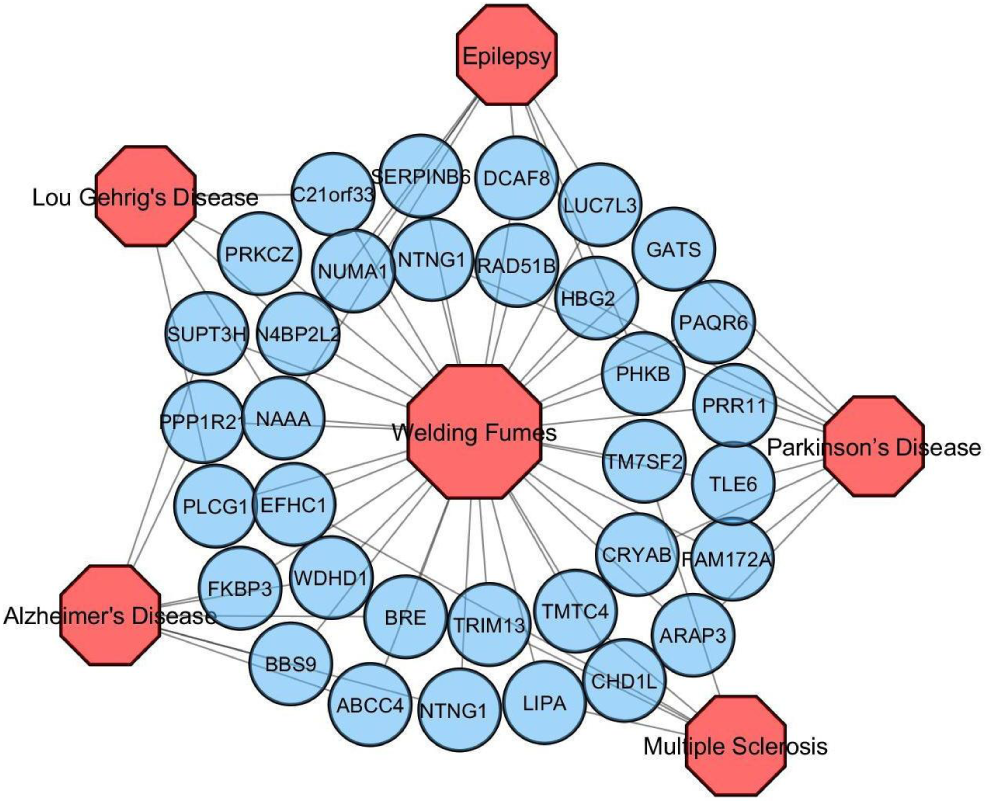
Disease network of Welding fumes (WFs) with Parkinson’s disease (PD), Alzheimer’s disease (AD), Lou Gehrig’s disease (LGD), Epilepsy disease (ED) and Multiple Sclerosis disease (MSD). Red colored octagon-shaped nodes represent different categories of disease, and round-shaped sky blue colored nodes represent commonly down-regulated genes for WFs with the other neurodegenerative disorders. A link is placed between a disorder and a disease gene if mutations in that gene lead to the specific disorder.

### 3.2 Pathway and Functional Association Analysis

Pathways are the key to know how an organism reacts to perturbations in its internal changes. The pathway-based analysis is a modern technique to understand how different complex diseases are related to each other by underlying molecular or biological mechanisms [24]. We analyzed pathways of the common differentially expressed genes using Enrichr, a comprehensive gene set enrichment analysis web-based tool [25]. Pathways of the commonly dysregulated genes in between WFs and each NDGD were analyzed using four databases includes KEGG, WikiPathways, Reactome and BioCarta. We combined pathways from four mentioned databases and identified the most significant pathways of each disease after various steps of statistical analysis.

We observed that PD has five significant pathways as shown in table 1. Among these pathways, ‘Glutamate Neurotransmitter Release Cycle’ is responsible to release the glutamate from the presynaptic neuron and its binding to glutamate receptors on the postsynaptic cell to generate a series of events that lead to the propagation of the synaptic transmission [26]. The pathway ‘Sphingolipid de novo biosynthesis’ is responsible to provide signals in molecules that regulate various biological functions [27]. The pathway ‘Intrinsic Pathway for Apoptosis’ is responsible to manage a variety of intracellular stress signal including DNA damage, growth factor withdrawal, unfolding stresses in the endoplasmic reticulum and death receptor stimulation [28]. Kinesins are a super-group of motor proteins based on microtubule that has various functions in the transport of vesicles, organelles, chromosomes, and regulate microtubule dynamics [29]. The pathway ‘Neurotransmitter Release Cycle’ is responsible to control electrical signals passing through the axons in the form of action potential.

**Table 1:** Pathways Associated with Significantly Commonly Differentially Expressed Genes of the PD with WFs.

| Pathway | Genes in the pathway | Adjusted p-value |
| --- | --- | --- |
| Glutamate Neurotransmitter Release Cycle | GLS | 2.02E-02 |
| Sphingolipid De Novo Biosynthesis | VAPB | 2.77E-02 |
| Intrinsic Pathway for Apoptosis | PMAIP1 | 3.51E-02 |
| Kinesins Pathway | KIF1A | 3.68E-02 |
| Neurotransmitter Release Cycle | GLS | 4.25E-02 |

We observed that AD has four significant pathways as shown in table 2. Among these pathways, ‘Circadian rhythm pathway’ is responsible to feed and influence clocks in other tissues by hormone secretion and nervous stimulation from the brain [30]. Sphingomyelin synthesis appears to be regulated primarily at the level of this transport process through the reversible phosphorylation of CERT (Saito et al. 2008). ‘Amyotrophic lateral sclerosis (ALS)’ is responsible for most common motor neuron disease [31]. ‘MAPKinase Signaling Pathway’ is responsible for manage signals of reactions that regulate cell proliferation and apoptosis [32].

**Table 2:** Pathways Associated with Significantly Commonly Differentially Expressed Genes of the AD with WFs.

| Pathway | Genes in the pathway | Adjusted p-value |
| --- | --- | --- |
| Circadian rhythm pathway | BHLHE40 | 2.23E-02 |
| Sphingolipid metabolism pathway | SGMS2 | 3.47E-02 |
| Amyotrophic lateral sclerosis (ALS) | MAP2K3 | 3.76E-02 |
| MAPKinase Signaling Pathway | MAP2K3 | 4.12E-02 |

We observed that LGD has six significant pathways as shown in table 3. Among these pathways, ‘Rap1 signaling pathway’ is responsible for controlling a variety of processes, such as cell adhesion, cell polarity and cell-cell junction formation [33]. ‘P53 signaling pathway’ manages various stress signals, including activated oncogenes, oxidative stress and DNA damage.

**Table 3:** Pathways Associated with Significantly Common Differentially Expressed Genes of the LGD with WFs.

| Pathway | Genes in the pathway | Adjusted p-value |
| --- | --- | --- |
| Signaling Pathways in Glioblastoma | CDKN1A, PLCG1, PRKCZ | 1.48E-05 |
| MAPK Signaling Pathway | CD14;PRKCZ | 4.38E-03 |
| TRIF-mediated programmed cell death | CD14 | 5.99E-03 |
| EPO Signaling Pathway | PLCG1 | 6.58E-03 |
| Rap1 signaling pathway | PLCG1, PRKCZ | 6.82E-03 |
| P53 signaling pathway | CDKN1A | 4.06E-02 |

We observed that ED has five significant pathways as shown in table 4. Among these pathways, ‘Neurotransmitter Release Cycle’ is responsible to control electrical signals passing through the axons in the form of action potential. ‘Glycogen Metabolismserves’ serves as a major stored fuel for several tissues. The keratinocytes function is to form a barrier against environmental damage by fungi pathogenic bacteria, parasites, viruses, and UV radiation.

**Table 4:**
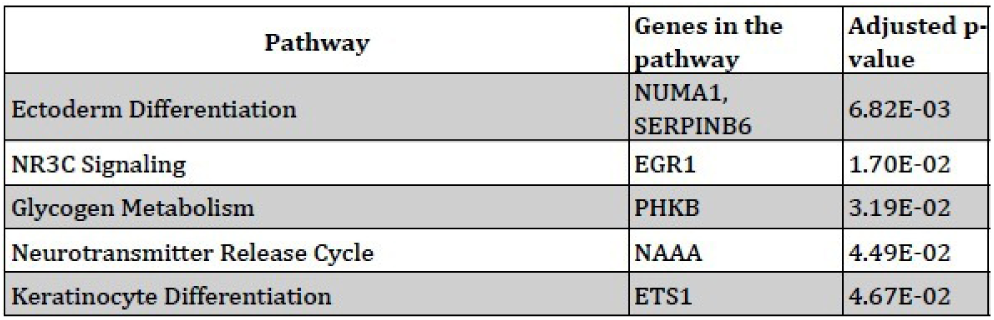
Pathways Associated with Significantly Common Differentially Expressed Genes of the ED with WFs.

| Pathway | Genes in the pathway | Adjusted p-value |
| --- | --- | --- |
| Ectoderm Differentiation | NUMA1, SERPINB6 | 6.82E-03 |
| NR3C Signaling | EGR1 | 1.70E-02 |
| Glycogen Metabolism | PHKB | 3.19E-02 |
| Neurotransmitter Release Cycle | NAAA | 4.49E-02 |
| Keratinocyte Differentiation | ETS1 | 4.67E-02 |

We observed that MSD has five significant pathways as shown in table 5. Among these pathways, ‘Endocrine and other factor-regulated calcium reabsorption’ is essential for numerous physiological functions including muscle contraction, intracellular signalling processes, neuronal excitability and bone formation [34]. ‘Mineral absorption’ provides mineral in the neural cell to sustain life. ‘Cholesterol biosynthesis’ controls cholesterol to the nucleus and activating genes.

**Table 5:** Pathways Associated with Significantly Common Differentially Expressed Genes of the MSD with WFs.

| Pathway | Genes in the pathway | Adjusted p-value |
| --- | --- | --- |
| Steroid biosynthesis | LIPA,TM7SF2 | 1.44E-04 |
| Metabolism of lipids and lipoproteins | CDS1, NCEH1, SEC24D, TM7SF2 | 2.47E-03 |
| Cholesterol biosynthesis | TM7SF2 | 2.05E-02 |
| Endocrine and other factor-regulated calcium reabsorption | ATP2B1 | 4.15E-02 |
| Mineral absorption | ATP2B1 | 4.49E-02 |

### 3.3 Gene Ontological Analysis

The Gene Ontology (GO) refers to a universal conceptual model for representing gene functions and their relationship in the domain of gene regulation. It is constantly expanded by accumulating the biological knowledge to cover regulation of gene functions and the relationship of these functions in terms of ontology classes and semantic relations between classes [35]. GO of the significantly dysregulated genes were analyzed using Enrichr, a comprehensive gene set enrichment analysis web-based tool [25]. GO of the commonly differentially expressed genes (i.e. Dysregulated genes in between WFs and each NDGD) for each NDGD and WFs were analyzed using two databases of Enrichr including GO Biological Process and Human Phenotype Ontology. We combined ontologies from two mentioned databases and identified the most significant GO term of each disease after various steps of statistical analysis. We observed that 15, 15, 24, 19 and 17 gene ontology classes are associated with the significantly commonly dysregulated (i.e. Dysregulated genes in between WFs and each NDGD) genes for WFs with the PD, AD, LGD, Ed and MSD respectively as shown in table 6-10.

**Table 6:** Gene Ontologies Associated with the Significantly Commonly Dysregulated Genes of the PD with WFs.

| GO Term | Pathway | Genes in the Pathway | Adjusted p-value |
| --- | --- | --- | --- |
| GO:0001844 | Protein insertion into mitochondrial membrane involved in apoptotic signaling pathway | PMAIP1 | 5.94E-03 |
| GO:1902043 | Positive regulation of extrinsic apoptotic signaling pathway via death domain receptors | PMAIP1 | 5.94E-03 |
| GO:0006987 | Activation of signaling protein activity involved in unfolded protein response | VAPB | 6.78E-03 |
| GO:0001881 | Receptor recycling | LMTK2 | 8.47E-03 |
| GO:0045837 | Negative regulation of membrane potential | PMAIP1 | 5.94E-03 |
| GO:0043029 | T cell homeostasis | PMAIP1 | 9.31E-03 |
| GO:0032463 | Negative regulation of protein homooligomerization | CRYAB | 7.63E-03 |
| GO:0032075 | Positive regulation of nuclease activity | VAPB | 9.31E-03 |
| GO:0016192 | Vesicle-mediated transport | LMTK2, ARAP3, KIF1A | 4.73E-03 |
| GO:1905906 | Regulation of amyloid fibril formation | CRYAB | 7.63E-03 |
| HP:0003677 | Slow progression | KIF1A, CRYAB | 3.08E-03 |
| HP:0007210 | Lower limb amyotrophy | KIF1A | 8.47E-03 |
| HP:0200073 | Respiratory insufficiency due to defective ciliary clearance | ZMYND10 | 7.63E-03 |
| HP:0003323 | Progressive muscle weakness | VAPB | 9.31E-03 |
| HP:0003555 | Muscle fiber splitting | CRYAB | 6.78E-03 |

### 3.4 Protein-Protein Interaction Analysis

Protein-protein interaction networks (PPINs) are the mathematical representation of the physical contacts of proteins in the cell. Protein-protein interactions (PPIs) are essential to every molecular and biological process in a cell, so PPIs is crucial to understand cell physiology in disease and healthy states [36]. PPIs of the differentially expressed genes were analyzed using STRING, a biological database and web resource of known and predicted protein-protein interactions [37]. We constructed protein-protein interaction network of significantly commonly dysregulated genes (i.e. Dysregulated genes in between WFs and each NDGD) of all NDGDs using STRING. We clustered into five different groups of interactions of five NDGDs as shown in figure 4.

**Figure 4:**
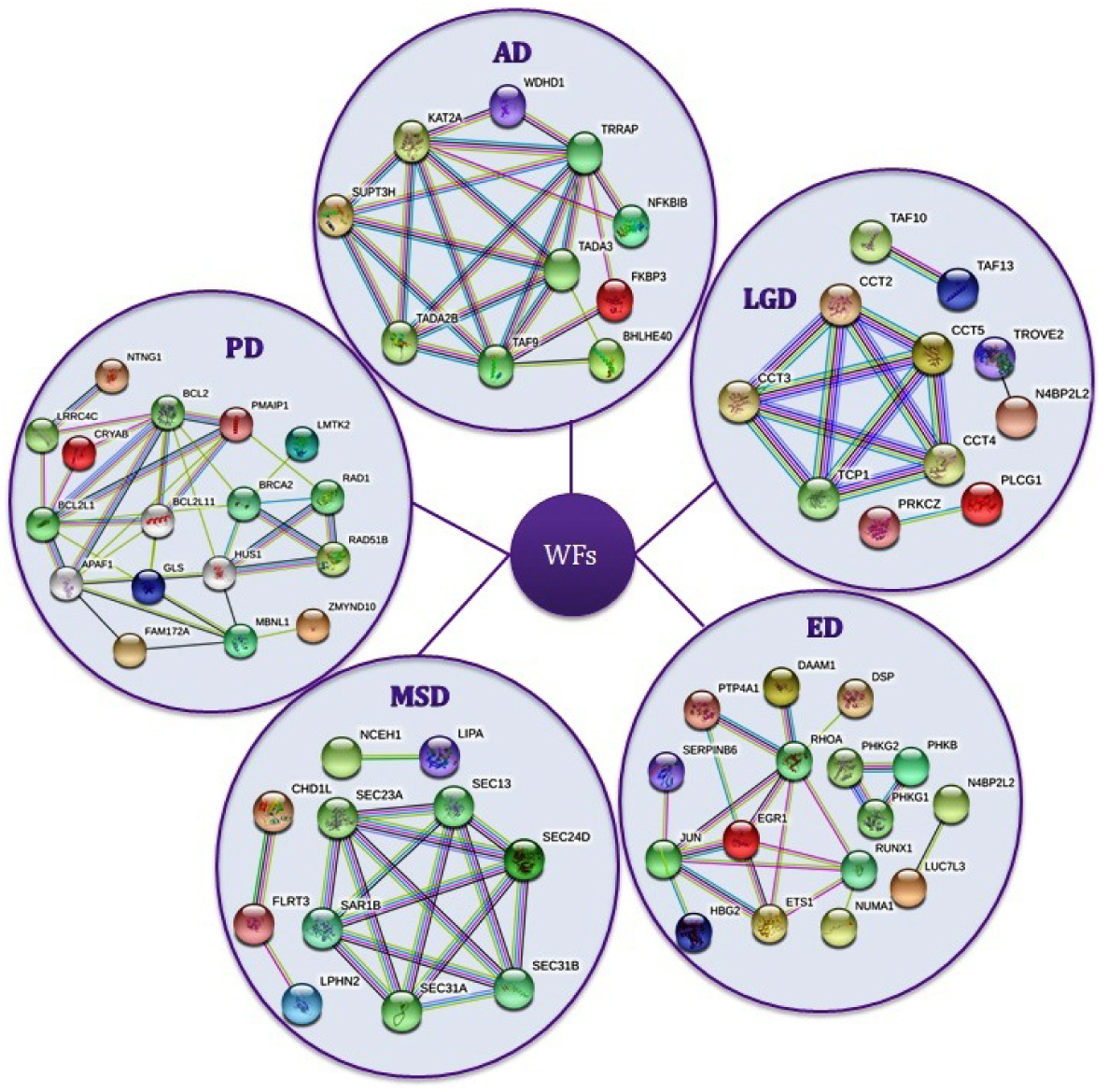
Protein-Protein Interaction Network of the Significantly Commonly Dysregulated Genes of the NDGDs with WFs.

## 4. Discussion

We investigated the genetic relationship of Welding fumes (WFs) and neurodegenerative diseases (NDGDs) based on the associations of genetics, signaling pathways, gene expression ontologies and protein-protein interactions network. For the purpose of our study, we analyzed Gene Expression Omnibus (GEO) microarray data from WFs, Parkinson’s disease (PD), Alzheimer’s disease (AD), Lou Gehrig’s disease (LGD), Epilepsy disease (ED), Multiple Sclerosis disease (MSD) and control datasets. We found a good number of significantly commonly dysregulated genes in between WFs and NDGDs by gene expression analysis. As there have a good number of significantly commonly dysregulated genes of WFs and NDGDs, it determines that WFs should have effects on NDGDs. Our two separate disease relationships networks for up and down-regulated genes strongly indicated that WFs are highly responsible for NDGDs as shown in Figure 2 and 3. The pathway-based analysis is a modern technique to understand how different complex diseases are related to each other by underlying molecular or biological mechanisms. We identified pathways of the commonly differentially expressed genes (i.e. Dysregulated genes in between WFs and each NDGD) of each NDGD. These identified pathways agreed that WFs have a strong association with NDGDs. Similarly, gene expression ontologies and protein-protein interactions of common differentially expressed genes determine that WFs can carry several NDGDs on unsafe welder’s health.

We have verified our result with the gold benchmark databases (dbGAP and OMIM) and found that there are some shared genes between the WFS and NDGDs as shown in figure 5. For cross checking the validity of our study, we collected genes and disease names from OMIM Disease, OMIM Expanded and dbGap databases using differentially expressed genes of WFs. We combined the diseases from three mentioned databases and selected only neurodegenerative diseases (NDGDs) after various steps of statistical analysis. Interestingly, we found our selected five NDGDs among the list of collected NDGDs from the mentioned databases as shown in figure 5. Therefore, it proved that WFs may have a strong association for the progression of PD, AD, LGD, ED and MSD neurodegenerative diseases [7, 13].

**Figure 5:**
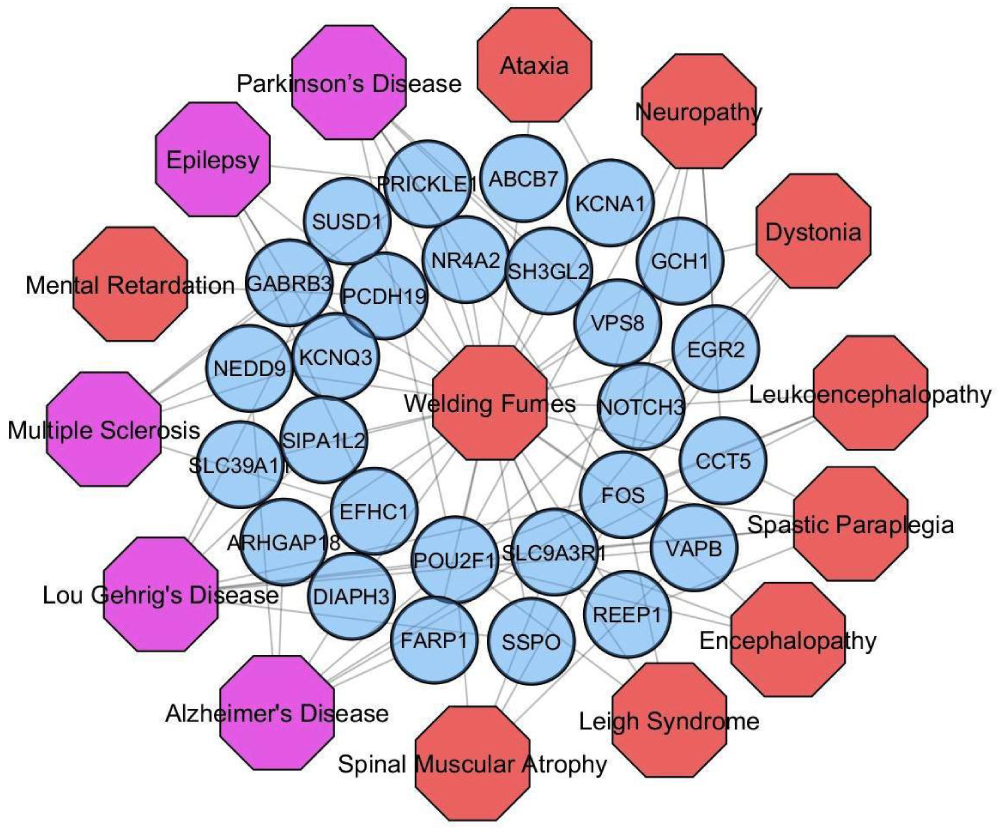
Disease network of Welding fumes (WFs) with several NDGDs. Red colored octagon-shaped nodes represent different categories of NDGDs, Violet colored octagon-shaped nodes represent our selected five NDGDs and round-shaped sky blue colored nodes represent differentially expressed genes for WFs. A link is placed between a disorder and a disease gene if mutations in that gene lead to the specific disorder

## 5. Conclusions

In this study, we have considered Gene Expression Omnibus (GEO) microarray data from welding fumes (WFs), Parkinson’s disease (PD), Alzheimer’s disease (AD), Lou Gehrig’s disease (LGD), Epilepsy disease (ED), Multiple Sclerosis disease (MSD) and control datasets to analyze and investigate the genetic effects of WFs on neurodegenerative diseases (NDGDs). We analyzed dysregulated genes, disease relationship networks, dysregulated pathways, gene expression ontologies and protein-protein interactions of WFs and NDGDs. Our findings showed that WFs have a strong association with NDGDs. This kind of study will be useful for making genomic evidence based recommendations about the accurate disease prediction, identification and therapeutic treatments. This study also will be useful for making society aware of the dangerous effect of welding on the human body.

**Table 7:** Gene Ontologies Associated with the Significantly Commonly Dysregulated Genes of the AD with WFs.

| GO Term | Pathway | Genes in the Pathway | Adjusted p-value |
| --- | --- | --- | --- |
| GO:0043619 | Regulation of transcription from RNA polymerase II promoter in response to oxidative stress | SESN2 | 6.73E-03 |
| GO:0032055 | Negative regulation of translation in response to stress | SESN2 | 5.24E-03 |
| GO:0006684 | Sphingomyelin metabolic process | SGMS2 | 7.48E-03 |
| GO:0035414 | Negative regulation of catenin import into nucleus | WWTR1 | 7.48E-03 |
| GO:1990253 | Cellular response to leucine starvation | SESN2 | 8.22E-03 |
| GO:0032309 | Icosanoid secretion | ABCC4 | 8.97E-03 |
| GO:0045859 | Regulation of protein kinase activity | MAP2K3, WWTR1 | 5.36E-03 |
| HP:0001156 | Brachydactyly syndrome | NTNG1, BBS9 | 8.87E-03 |
| HP:0002141 | Gait imbalance | BBS9 | 1.34E-02 |
| HP:0007707 | Congenital primary aphakia | BBS9 | 1.34E-02 |
| HP:0010747 | Medial flaring of the eyebrow | BBS9 | 1.56E-02 |
| HP:0009806 | Nephrogenic diabetes insipidus | BBS9 | 1.49E-02 |
| HP:0002370 | Poor coordination | BBS9 | 1.71E-02 |
| HP:0006829 | Severe muscular hypotonia | IBA57 | 1.79E-02 |
| HP:0001827 | Genital tract atresia | BBS9 | 1.93E-02 |

**Table 8:** Gene Ontologies Associated with the Significantly Commonly Dysregulated Genes of the LGD with WFs.

| GO Term | Pathway | Genes in the Pathway | Adjusted p-value |
| --- | --- | --- | --- |
| GO:2000737 | Negative regulation of stem cell differentiation | N4BP2L2, ZFP36L2 | 1.81E-05 |
| GO:0034128 | Negative regulation of MyD88-independent toll-like receptor signaling pathway | CD14 | 4.79E-03 |
| GO:0071364 | Cellular response to epidermal growth factor stimulus | PLCG1, ZFP36L2 | 1.06E-04 |
| GO:1901988 | Negative regulation of cell cycle phase transition | ZFP36L2 | 4.19E-03 |
| GO:1903708 | Positive regulation of hemopoiesis | N4BP2L2 | 4.79E-03 |
| GO:1901991 | Negative regulation of mitotic cell cycle phase transition | CDKN1A, ZFP36L2 | 1.02E-03 |
| GO:0071363 | Cellular response to growth factor stimulus | PLCG1, ZFP36L2 | 3.07E-03 |
| GO:0050821 | Protein stabilization | CDKN1A, CCT5 | 3.65E-03 |
| HP:0001738 | Exocrine pancreatic insufficiency | CDKN1A | 1.13E-02 |
| HP:0010832 | Abnormality of pain sensation | CCT5 | 1.49E-02 |
| HP:0002717 | Adrenal overactivity | CDKN1A | 1.79E-02 |
| HP:0003431 | Decreased motor nerve conduction velocity | CCT5 | 1.79E-02 |
| HP:0001258 | Spastic paraplegia | CCT5 | 1.84E-02 |
| HP:0002936 | Distal sensory impairment | CCT5 | 3.66E-02 |

**Table 9:**
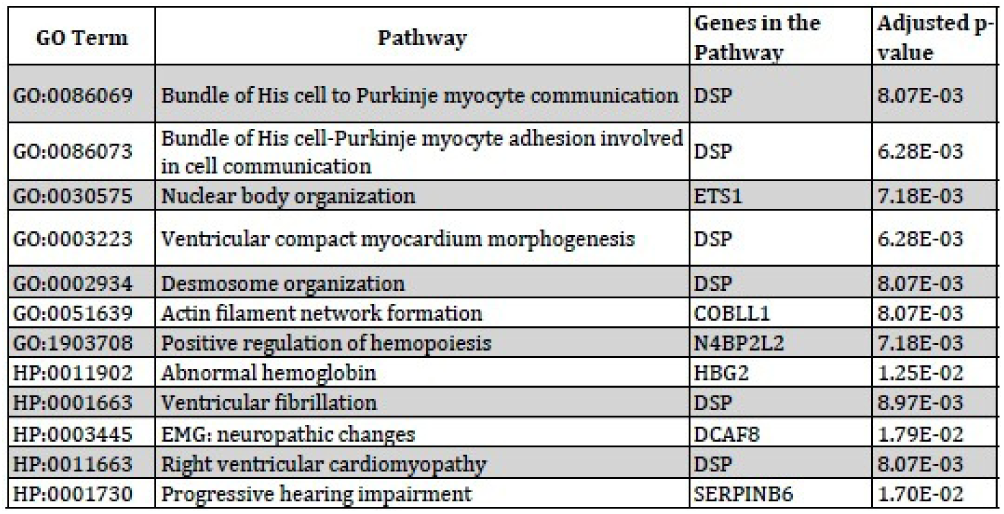
Gene Ontologies Associated with the Significantly Commonly Dysregulated Genes of the ED with WFs.

| GO Term | Pathway | Genes in the Pathway | Adjusted p-value |
| --- | --- | --- | --- |
| GO:0086069 | Bundle of His cell to Purkinje myocyte communication | DSP | 8.07E-03 |
| GO:0086073 | Bundle of His cell-Purkinje myocyte adhesion involved in cell communication | DSP | 6.28E-03 |
| GO:0030575 | Nuclear body organization | ETS1 | 7.18E-03 |
| GO:0003223 | Ventricular compact myocardium morphogenesis | DSP | 6.28E-03 |
| GO:0002934 | Desmosome organization | DSP | 8.07E-03 |
| GO:0051639 | Actin filament network formation | COBLL1 | 8.07E-03 |
| GO:1903708 | Positive regulation of hemopoiesis | N4BP2L2 | 7.18E-03 |
| HP:0011902 | Abnormal hemoglobin | HBG2 | 1.25E-02 |
| HP:0001663 | Ventricular fibrillation | DSP | 8.97E-03 |
| HP:0003445 | EMG: neuropathic changes | DCAF8 | 1.79E-02 |
| HP:0011663 | Right ventricular cardiomyopathy | DSP | 8.07E-03 |
| HP:0001730 | Progressive hearing impairment | SERPINB6 | 1.70E-02 |

**Table 10:** Gene Ontologies Associated with the Significantly Commonly Dysregulated Genes of the MSD with WFs.

| GO Term | Pathway | Genes in the Pathway | Adjusted p-value |
| --- | --- | --- | --- |
| GO:0021795 | Cerebral cortex cell migration | EFHC1 | 6.28E-03 |
| GO:0048678 | Response to axon injury | FLRT3 | 1.07E-02 |
| GO:1902187 | Negative regulation of viral release from host cell | TRIM13 | 1.25E-02 |
| GO:0014033 | Neural crest cell differentiation | KBTBD8 | 1.43E-02 |
| GO:0006293 | Nucleotide-excision repair, preincision complex stabilization | CHD1L | 1.96E-02 |
| HP:0004311 | Abnormality of macrophages | LIPA | 1.70E-02 |
| HP:0001433 | Hepatosplenomegaly | LIPA | 2.23E-02 |
| HP:0100639 | Erectile abnormalities | FLRT3 | 2.84E-02 |
| HP:0002612 | Congenital hepatic fibrosis | LIPA | 3.37E-02 |
| HP:0001522 | Death in infancy | LIPA | 3.98E-02 |

